# Illumina But With Nanopore: Sequencing Illumina libraries at high accuracy on the ONT MinION using R2C2

**DOI:** 10.1101/2021.10.30.466545

**Authors:** Alexander Zee, Dori Z. Q. Deng, Matthew Adams, Kayla D. Schimke, Russell Corbett-Detig, Shelbi L. Russell, Xuan Zhang, Robert J. Schmitz, Christopher Vollmers

## Abstract

High-throughput short-read sequencing has taken on a central role in research and diagnostics. Hundreds of different assays exist today to take advantage of Illumina short-read sequencers, the predominant short-read sequencing technology available today. Although other short read sequencing technologies exist, the ubiquity of Illumina sequencers in sequencing core facilities, and the high capital costs of these technologies have limited their adoption. Among a new generation of sequencing technologies, Oxford Nanopore Technologies (ONT) holds a unique position because the ONT MinION, an error-prone long-read sequencer, is associated with little to no capital cost. Here we show that we can make short-read Illumina libraries compatible with the ONT MinION by using the R2C2 method to circularize and amplify the short library molecules. This results in longer DNA molecules containing tandem repeats of the original short library molecules. This longer DNA is ideally suited for the ONT MinION, and after sequencing, the tandem repeats in the resulting raw reads can be converted into high-accuracy consensus reads with similar error rates to that of the Illumina MiSeq. We highlight this capability by producing and benchmarking RNA-seq, ChIP-seq, as well as regular and target-enriched Tn5 libraries. We also explore the use of this approach for rapid evaluation of sequencing library metrics by implementing a real-time analysis workflow.

## Introduction

Over the last 15 years, high-throughput short-read sequencing technology has revolutionized biological, biomedical, and clinical research. Hundreds of sequencing based methods exist today to query gene expression (RNA-seq(1)), chromatin state (ChIP-seq(2) and ATAC-seq(3)), protein abundance(4), and of course to aid the assembly of genomes(5) - among many other things. All of these methods produce a final sequencing library that contains ~200-600bp double stranded DNA molecules with ends of a known sequence. In the vast majority of cases, these ends are Illumina sequencing adapters.

Despite the existence of other sequencing technologies, Illumina has been the dominating short-read sequencing technology over the last decade. However, due to the high capital cost of Illumina short-read instruments, all but the most well equipped labs outsource their Illumina sequencing to core facilities. While this provides access to the most recent sequencing technology, this outsourcing can lead to long delays between running an experiment and receiving results. Therefore, placing a benchtop sequencer with capabilities comparable to an Illumina sequencer in most molecular biology and diagnostic labs could be truly transformative by accelerating as well as fully integrating genomics assays into standard lab workflows. In a molecular biology lab, it would speed up developing or establishing new types of sequencing libraries. In a diagnostic lab it could enable fast sample turn-around as well as encourage the transition away from outdated and flawed diagnostic methods like Fluorescence In Situ Hybridization (FISH).

Over the last few years Oxford Nanopore Technologies (ONT) sequencers have rapidly matured. Currently, the ONT MinION sequencer’s base throughput (up to 30 Gb per flow cell) can exceed that of the Illumina MiSeq sequencer (18 Gb for a 2×300 bp run). Intriguingly, this throughput comes with tunable read length, so a succesfull MinION run can in theory produce 10 million 3kb reads or 5 million 6kb reads. Further, the MinION sequencer is only a fraction of the cost of other high-throughput sequencers.

However, standard per-base sequencing accuracy of the newest basecalling software guppy5 is only around 96% and dominated by insertion and deletion errors which are almost absent in Illumina data. Furthermore, ONT MinION’s sequencing accuracy declines with shorter reads(8).

Here, we implemented a simple workflow that converts almost any Illumina sequencing library into DNA of lengths optimal for the ONT MinION and generates data at similar cost and accuracy as the Illumina MiSeq. We made this possible by using the previously published and optimized R2C2 (<u>R</u>olling Circle <u>to C</u>oncatemeric <u>C</u>onsensus) method(9–14). R2C2 circularizes dsDNA libraries and amplifies those circles using rolling circle amplification to create long molecules with multiple tandem repeats of the original molecule’s sequence. These long molecules can then be sequenced on ONT instruments to generate long raw reads which are then computationally processed into accurate consensus reads. In previous studies focused on full-length cDNA molecules we have achieved median read accuracies of 99.5% with this method(13). Since Illumina libraries are shorter than full-length cDNA, we modified the R2C2 protocol to generate a large number of shorter MinION raw reads while maintaining consensus accuracy levels on par with the Illumina MiSeq sequencer.

We benchmark this extension of the R2C2 method by converting and sequencing RNA-seq, ChIP-seq, as well as regular and target-enriched genomic DNA Tn5 Illumina libraries. We implemented a computational workflow for demultiplexing Illumina library indexes from R2C2 data and have, where possible, relied on established analysis workflows for downstream analysis. To highlight the potential of this approach for the rapid evaluation of library metrics, we developed the PLNK (<u>P</u>rocessing <u>L</u>ive <u>N</u>anopore Experiments) tool to take advantage of the real-time data generation of the ONT MinION. PLNK processes raw data and generates immediate feedback on library composition by reporting the percentage of reads that map to target regions in the genome as well as the coverage over these regions. We show that this on-target percentage can be used to evaluate RNA-seq, ChIP-seq and enriched Tn5 libraries.

## Results

To generate R2C2 data for a diverse selection of Illumina libraries, we processed and sequenced 1) Illumina RNA-seq libraries of the human A549 cancer cell line, 2) Illumina ChIP-seq and Input libraries of soybean samples, 3) Illumina Tn5-based genomic DNA libraries of a *Wolbachia*-containing *Drosophila melanogaster* cell line, and 4) Illumina Tn5-based genomic DNA libraries generated from lung cancer cell lines NCI-H1650 and NCI-H1975 which we enriched for the protein coding regions of ~100 cancer relevant genes (Fig. 1).

**Fig. 1:**
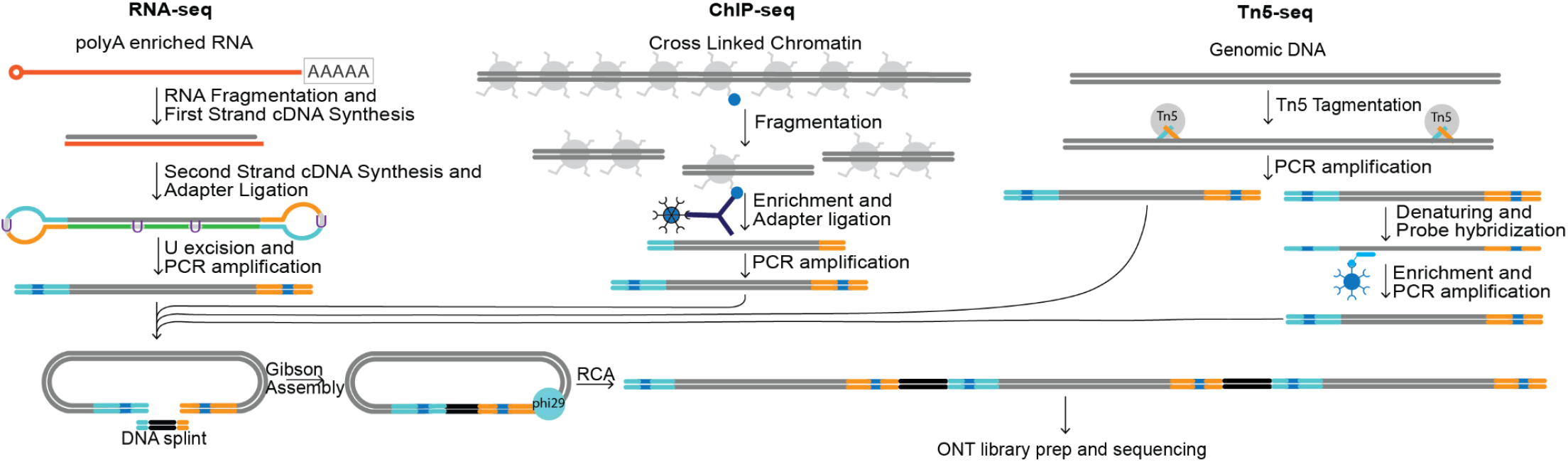
Experiment overview. Illumina RNA-seq, ChIP-seq, and Tn5-based genomic libraries libraries (regular and enriched) were generated from different samples. The Illumina libraries were then circularized and amplified using rolling circle amplification (RCA). The resulting DNA, containing tandem repeats of Illumina library molecules, was then prepped for sequencing on the ONT MinION sequencer.

To convert these Illumina libraries into R2C2 libraries, we circularized them using Gibson assembly (NEBuilder/NEB) with DNA splints compatible with Illumina p5 and p7 sequences (Table S1). After the DNA circles are amplified with rolling circle amplification using Phi29 polymerase, we fragmented and size selected the resulting high molecular weight DNA. We then sequenced this DNA on the ONT MinION using the LSK-110 ligation chemistry and 9.4.1 flow cells. We generated between 4 and 9.5 million raw reads per MinION flow cell (Table 1). All data was then basecalled with the guppy5 *dna_r9.4.1_450bps_sup.cfg* model and consensus called using C3POa (v2.2.3) (https://github.com/rvolden/C3POa).

**Table 1:** R2C2 sequencing run characteristics. For consistency, median per read accuracy is calculated for R2C2 reads prior to demultiplexing.

| Library type | Organism | Raw reads (pass filter) | Raw read median length | R2C2 reads | Demuxed reads | Subreads/ R2C2 read | Median Per-Read accuracy |
| --- | --- | --- | --- | --- | --- | --- | --- |
| RNA-seq | Homo sapiens | 9,500,956 | 2,288 | 8,992,882 | 8,066,704 | 3.14 | 99.52% |
| ChIP-seq | Glycine max (soybean) | 4,518,775 | 3,360 | 4,191,438 | 4,023,935 | 3.93 | 99.12% |
| Tn5 | D. melanogaster/ Wolbachia | 5,188,771 | 2,447 | 3,339,161 | N/A | 4.88/ 4.68 | 98.8%/ 99.61% |
| Enriched Tn5 | Homo sapiens | 4,062,736 | 3,377 | 3,825,657 | 3,078,913 | 4.85 | 99.38% |

To benchmark the R2C2 data for the Illumina libraries, we sequenced the same libraries with regular ONT 1D reads and on different Illumina sequencers. We then compared the metrics most relevant to the different library types.

### Evaluating R2C2 for the sequencing of Illumina RNA-seq libraries

First, we benchmarked the ONT-based R2C2 method for the generation of RNA-seq data from Illumina libraries. We prepared four technical replicates in the form of dual indexed paired-end Illumina libraries using the NEBnext Ultra II Directional RNA kit with the human lung carcinoma cell line A549. We pooled and sequenced these libraries with the ONT MinION both directly (1D) and after R2C2 conversion (R2C2) as well as with the Illumina MiSeq.

To establish the effect of R2C2 conversion on the throughput of the ONT MinION when sequencing short Illumina libraries, we processed the raw reads generated by both 1D and R2C2 sequencing runs. Raw read numbers for 1D and R2C2 runs generated from one ONT MinION flow cell were similar at ~11.8 million reads. However, 1D reads were less likely than R2C2 reads to 1) pass filter during basecalling, 2) contain both p5 and p7 Illumina adapter sequences, and 3) be successfully demultiplexed. After preprocessing, only 2.5 million 1D reads remained compared to ~8 million R2C2 reads (Table 2). This means that even a much more productive 1D run, potentially generating up to 20 million raw reads for molecules of this length (15), would still generate fewer demultiplexed reads (~5 million) than the R2C2 run we performed here.

**Table 2.** R2C2 and 1D read numbers throughout processing steps

| Library type | Raw reads | pass filter | Consensus reads | p5/p7 adapters present | demultiplexed |
| --- | --- | --- | --- | --- | --- |
| R2C2 (ONT MinION) | 11,789,059 | 9,500,956 | 9,132,280 | 8,992,882 | 8,066,704 |
| 1D (ONT MinION) | 11,839,886 | 7,578,968 | N/A | 3,469,357 | 2,530,950 |

To validate the demultiplexing of Illumina library pools from R2C2 data, we compared the ratio of reads assigned to each library in Illumina MiSeq, R2C2, and ONT 1D data. For all three methods, three technical replicate libraries were pooled at 4:2:1 ratio. The Illumina MiSeq produced 4:2.03:1.58 read ratio after demultiplexing. R2C2 produced a 4:1.91:1.34 ratio and ONT 1D produced a 4:2.5:1.82 ratio. With these results being quite similar, the differences are likely due to pipetting variability when pooling the libraries for the different sequencing methods. Further, to evaluate our ability to quantitatively pool libraries at different points in the R2C2 workflow, we processed a fourth replicate in parallel and added it at a specific ratio after rolling circle amplification. The fourth replicate represented 40.5% of the R2C2 data which is slightly more than the 30% of R2C2 DNA it represented in the MinION sequencing run. Finally, while 9.71% of R2C2 reads were not assigned to any sample, only 1.7% of R2C2 reads were assigned to a combination of Illumina indexes not included in the pool. The 1.7% single index switching rate implies that theoretically only 0.0289% (1.7%*1.7%) or less than 3 out of 10,000 reads were misassigned to the wrong sample in a dual indexed library.

Next we established the effect of R2C2 conversion on read accuracy when compared to ONT 1D and Illumina MiSeq datasets. We aligned all complete p5 and p7 containing and demultiplexed R2C2 (8,066,704) and 1D reads (2,530,950) as well as Illumina MiSeq reads (20,830,560 2×300 bp paired-end reads) generated from these RNA-seq libraries using minimap2. We then calculated the median read accuracy, accuracy per base, and read position dependent accuracy per base (Table 3).

**Table 3.** Sequencing error rates of different methods based on minimap2 alignments of all demultiplex reads

| Sequencing method | Median read accuracy (%) | Accuracy per base (%) | Mismatch rate per base (%) | Insertion rate per base (%) | Deletion rate per base (%) |
| --- | --- | --- | --- | --- | --- |
| R2C2 (ONT MinION) | 99.56 | 98.9 | 0.31 | 0.26 | 0.55 |
| 1D (ONT MinION) | 97.2 | 96.59 | 1.16 | 0.81 | 1.44 |
| Read 1 (Illumina MiSeq) | 100 | 99.47 | 0.45 | 0.04 | 0.04 |
| Read 2 (Illumina MiSeq) | 99.54 | 98.57 | 1.33 | 0.05 | 0.05 |

**Table 4.** ChIP-seq read characteristics

| Sample | Illumina HiSeq Reads | Median Insert length | R2C2 Reads | Median Insert length |
| --- | --- | --- | --- | --- |
| H3K4me3 | 8,413,865 | 389 | 2,493,021 | 556 |
| Input | 32,377,813 | 312 | 1,530,914 | 459 |

While median read accuracy is a useful and often reported metric to compare error-prone long-read sequencing technologies, it becomes less useful in this study. The sequencing reads we aim to compare are very short - either due to the short length of the molecules sequenced (1D and R2C2)(Fig. 2A) or technology limitations (Illumina MiSeq) - and often accurate enough to be unlikely to contain errors at that length, causing many individual sequencing reads to be 100% accurate. This is obvious with read 1 of the Illumina MiSeq having a median accuracy of 100% which contains little information on the real Illumina MiSeq error rate. Accuracy per base (%), i.e. (correct bases of all reads/all bases of all reads)*100, is a more useful metric to compare accurate short reads. Using this metric we see that 1D reads are the least accurate with an accuracy per base of 96.49%. R2C2 falls between Illumina MiSeq read 1 (99.47%) and read 2 (98.57%) with an accuracy per base of 98.68%. Interestingly, while R2C2 reads contained more deletion and insertion errors, they contain fewer mismatch errors than both Illumina MiSeq read 1 and read 2.

**Fig. 2.**
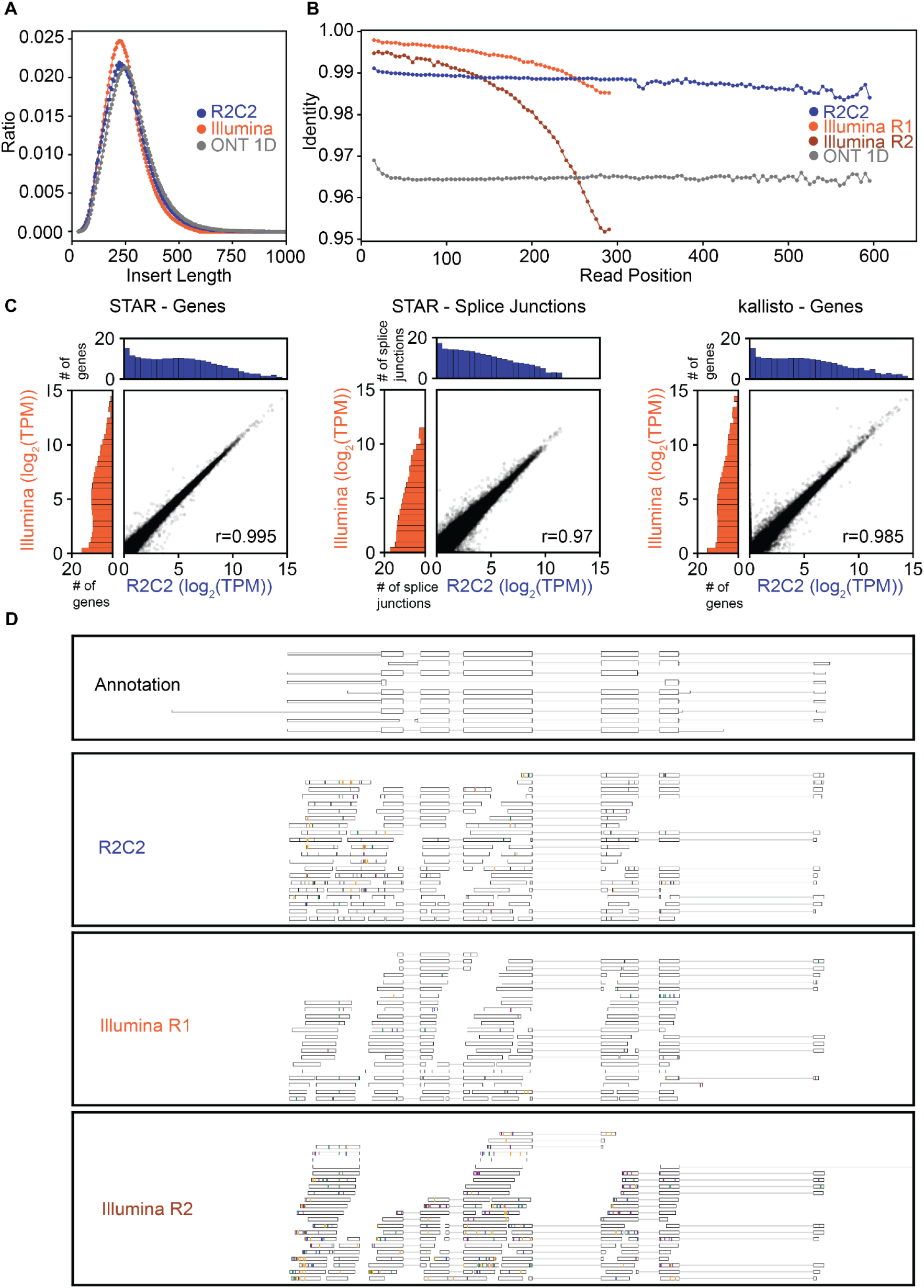
Sequencing Illumina RNA-seq libraries on the ONT MinION after R2C2 conversion. Insert length distribution (A) and read position dependent identity to the reference genome (B) of R2C2 and Illumina MiSeq reads generated from the same Illumina library. C) Comparisons of R2C2 and Illumina MiSeq read-based gene expression and splice junction usage quantification by STAR and kallisto are shown as scatter plots with marginal distributions (log2 normalized) shown as histograms. D) Genome browser-style visualization of read alignments. Mismatches are marked by lines colored by the read base (A - orange; T - green; C - blue; G - purple). Insertions are shown as gaps in the alignments while deletions are shown as black lines.

Read position dependent accuracy of 1D, R2C2, and Illumina MiSeq read 1 and read 2 adds further detail to this comparison. In contrast to 1D and R2C2 data, Illumina MiSeq base accuracy decreased with increasing read cycles, particularly in read 2, with R2C2 surpassing Illumina MiSeq accuracy for read 2 lengths over ~175 bp (Figure 2B and D).

Next, we aimed to establish whether R2C2 RNA-seq and ONT 1D data could be analyzed using computational tools designed and established for Illumina RNA-seq data. To quantify gene expression levels, we aligned and evaluated the entire demultiplexed R2C2 (8,066,704 reads) and ONT 1D (2,530,950 reads) datasets as well as our Illumina MiSeq dataset (20,830,560 read pairs) using the STAR aligner(16) (STARlong executable for R2C2 and ONT1D data) which is routinely used for standard Illumina RNA-seq analysis. 7,365,398 R2C2 reads (91.66%), 1,834,065 ONT 1D reads (72.48%) and 18,649,031 Illumina MiSeq reads (90.08%) mapped uniquely to the human genome, indicating that a larger percentage of ONT 1D reads aren’t accurate enough to be aligned by the STAR aligner.

Based on these read alignments, STAR determined normalized gene counts for Illumina MiSeq, R2C2, and ONT 1D datasets. Illumina MiSeq gene counts showed Pearson’s r-values of 0.995 and 0.987 when compared to R2C2 and ONT 1D, respectively (Fig. 2C). Additionally, STAR also determined normalized splice junction counts for the three datasets which provide a higher resolution view of the transcriptome. Illumina MiSeq splice junction counts showed Pearson’s r-values of 0.974 and 0.929 when compared to R2C2 and ONT 1D, respectively (Fig. 2C). Finally, we also tested whether ultra-fast pseudo-alignment based tools will generate reliable gene expression levels based on R2C2 and ONT 1D reads which feature more insertion and deletion rates compared to standard Illumina data. We used one such tool, kallisto (17), and found that gene expression values as determined for Illumina MiSeq had Pearson’s r values of 0.985 and 0.973 when compared to R2C2 and ONT 1D, respectively(Fig. 2C).

Overall this comparison showed that using R2C2, we can convert Illumina RNA-seq libraries into DNA ideally suited for the ONT MinION. Not only does R2C2 generate more reads than regular ONT 1D ligation protocols but R2C2 reads are also much more accurate. Because they are more accurate, R2C2 reads are also more efficiently demultiplexed and aligned than ONT 1D reads. Further, because they are similar in accuracy to Illumina reads, standard Illumina tools, like STAR and kallisto, can be used to analyze them. The gene expression and splice junction values generated by R2C2 are highly similar to those generated by Illumina MiSeq data from the same libraries.

### Evaluating R2C2 for the sequencing of Illumina ChIP-seq libraries

Next, we tested the ability of R2C2 for the quality control of Illumina ChIP-seq libraries. To do this, we converted a previously generated ChIP-seq library targeting the H3K4me3 histone modification in a *Glycine max* (soybean) sample. The H3K4me3 library and its corresponding control Input library had previously been sequenced on an Illumina NovaSeq 6000 to a depth of 8,413,865 and 32,377,813 2×150bp paired end reads, respectively. Based on their alignment, the sequenced molecule libraries had an insert length of 390 bp (H3K4me3) and 312 bp (Input).

Because the H3K4me3 and Input libraries were prepared with only a single index distinguishing them, we converted the libraries separately with R2C2 using distinct DNA splints that contained unique index sequences. This added an extra level of indexing to minimize concerns of potential index crosstalk. We splint-indexed, and pooled the H3K4me3 and Input ChIP-seq Illumina libraries and sequenced the pool on a single ONT MinION flow cell. We then demultiplexed the resulting R2C2 reads, assigning 2,493,021 and 1,530,914 reads (1.6:1) to the H3K4me3 and Input libraries, respectively, which a ratio which corresponded well with the 1.35:1 ratio at which they were pooled prior to sequencing. Importantly, the demultiplexing script scored only 163,489 (3.9%) reads as “undetermined” and assigned only 4,014 (0.1%) reads to a combination of indexes not present in the library. This indicated that the extra level of indexing was highly successful in minimizing index crosstalk.

The demultiplexed R2C2 reads showed a median read accuracy of 99.23% (H3K4me3) and 98.8% (Input) as well as a median read length of 556 bp (H3K4me3) and 459 bp (Input). The molecules sequenced by R2C2 were therefore longer than the molecules sequenced by the Illumina NovaSeq 6000. The difference between the technologies is likely due to the high bias of the Illumina NovaSeq 6000 towards shorter molecules.

To test whether R2C2 reads could replace the same number of Illumina reads, we subsampled the Illumina sequencing data to the depth of the R2C2 data for both samples. We then aligned both Illumina NovaSeq 6000, subsampled Illumina NovaSeq 6000, and R2C2 reads to the *Glycine max* genome (Gmax_508_v4.0)(17). For alignment, we chose the short-read preset of the minimap2(18) aligner for both Illumina and R2C2 data. We then called peaks on the full H3K4me3 Illumina NovaSeq 6000 dataset using MACS2 and tested whether both subsampled Illumina NovaSeq 6000 and R2C2 data could be used to evaluate the success of a ChIP experiment. Visual inspection of the data using the Phytozome JBrowse genome browser(19) as well as our own tools (Fig. 3D) showed that subsampled Illumina NovaSeq 6000 and R2C2 data both demonstrate the same enrichment patterns as the full Illumina NovaSeq 6000 data. A systematic analysis showed that 84% of R2C2 reads and 69% of subsampled Illlumina reads overlap with an H3K4me3 peak identified on the full Illumina data, whereas only 18% and 11% of the respective Input reads do so.

**Fig. 3.**
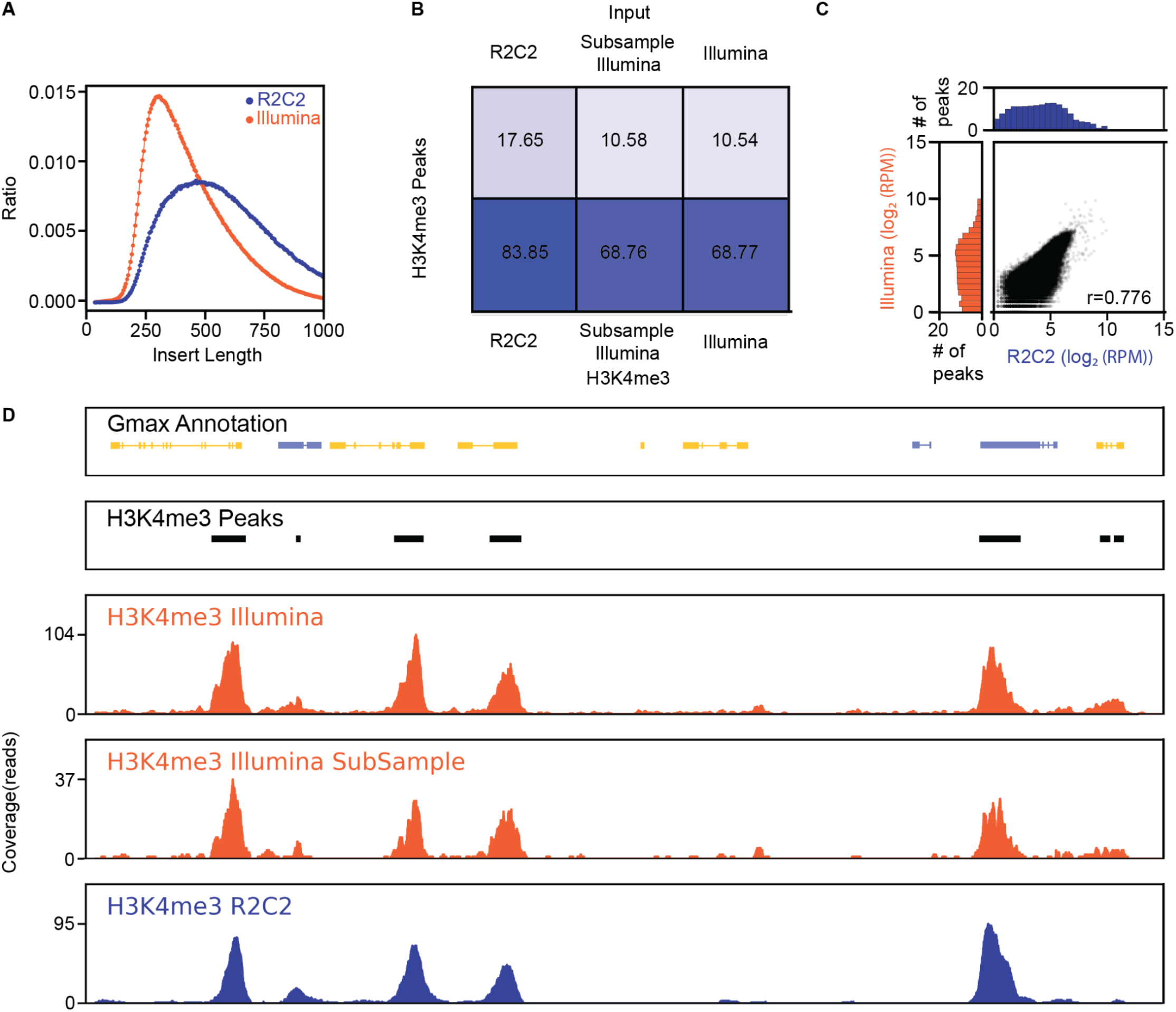
Sequencing Chip-seq libraries on the ONT MinION after R2C2 conversion. A) Insert length distribution of R2C2 and Illumina NovaSeq 6000 reads generated from the same Illumina library. B) Percentage of reads in the R2C2, Subsampled Illumina and full Illumina datasets overlapping with H3K4me3 peaks generated from the full Illumina H3K4me3 dataset using MACS2. C) Comparison of the number of R2C2 and subsampled Illumina reads overlapping with H3K4me3 peaks is shown as scatter plots with marginal distributions shown as histograms. Pearson’s r is shown in the bottom right. D) Genome annotation, H3K4me3 peak areas and read coverage histograms are shown for a section of the Gmax genome.

**Fig. 4.**
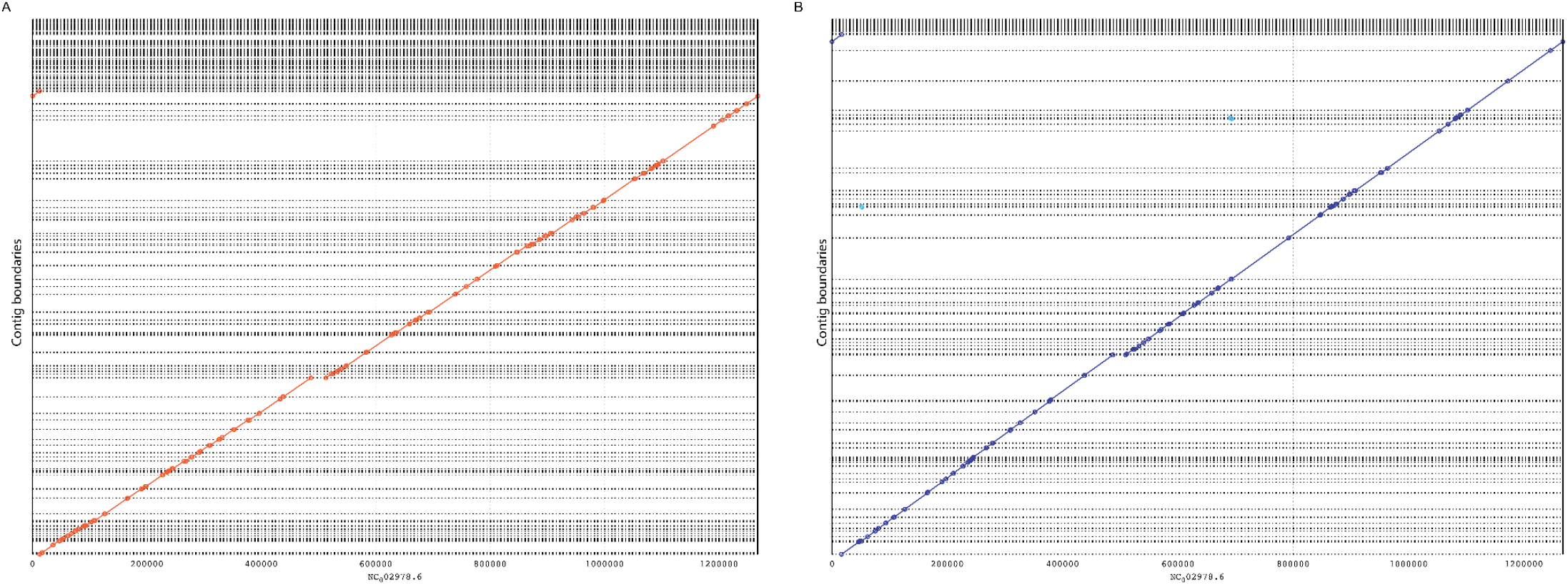
Comparing R2C2 and Illumina based assemblies of a small genome. Illumina 2×150 reads were assembled in 134 contigs using Meraculous. R2C2 reads were assembled using Miniasm into 95 contigs which were polished using Medaka. The alignments of the contigs of both assemblies - (A) Illumina and (B) R2C2 - are shown as dotplots generated by mummer(23). Both approaches fail to assemble a section of the Wolbachia genome that contains pseudogenes and a transposable element near to coordinate 500,000.

To compare whether R2C2 and subsampled Illumina NovaSeq 6000 datasets are also similar quantitatively, we counted how many reads for each of the datasets fell into each H3K4me3 peak we identified using the full Illumina NovaSeq 6000 dataset and MACS2. We found that the peak depths are correlated (Pearson’s r=0.776). This correlation is lower than we observed with the RNA-seq data, likely because the ChIP-seq library had a longer insert size than the RNA-seq library. This means that while R2C2 can be used to evaluate whether a ChIP-seq experiment successfully enriched chromatin from expected genomic regions, R2C2 will not produce quantification highly similar to the Illumina NovaSeq 6000.

### Evaluating R2C2 for the sequencing of Illumina Tn5 libraries

We tested whether R2C2 could be used to assemble small genomes from Illumina libraries generated using Tn5-based tagmentation. For this test, we chose the 1.2 Mb genome of the *Wolbachia* bacterial endosymbiont of *Drosophila melanogaster* and prepared Tn5 libraries from DNA extracted from *Wolbachia*-containing *Drosophila melanogaster* S2 cells.

While longer molecules are beneficial for genome assembly, Illumina sequencers struggle to generate clusters and sequence molecules longer than 600 bp. However, Tn5 generates molecules longer than this limit and R2C2 can efficiently process and sequence them. Therefore, instead of R2C2 converting the exact same library we used for Illumina sequencing, we chose to size-select an Illumina library. We size-selected a Tn5 library for molecules between 800-1200 bp lengths, corresponding to genomic DNA inserts of ~600-1000 bp. We then R2C2 converted and sequenced this size-selected Illumina library on the ONT MinION. We generated a total of 3,338,280 R2C2 consensus reads with a median length of 680 bp. Out of these reads, we assembled 879,303 reads that did not align to the *Drosophila melanogaster* genome. We used miniasm(20) for this assembly task and polished the resulting assembly using Medaka (v.1.4.4; https://github.com/nanoporetech/medaka). The resulting assembly contained 95 contigs which covered 97.2% of the *Wolbachia* genome, had a NGA50 of 29,963 bp and 8.5/5.6 mismatches/indels per 100 kb of sequence.

We also generated an assembly from Illumina Nextseq 2×150 bp generated from a non-size selected Tn5 library of the same cell line. From 2,552,018 2×150 bp Illumina reads we extracted 779,206 reads that did not align to the *Drosophila melanogaster* genome and assembled those reads using Meraculous(21). The resulting assembly contained 136 contigs which covered 91.6% of the *Wolbachia* genome, had a NGA50 of 23,217 bp and 0.5/0.6 mismatches/indels per 100 kb of sequence. Neither assembly had misassemblies as determined by QUAST(22).

Comparing Illumina and R2C2 assemblies of the *Wolbachia* genome (NC_002978.6) showed R2C2 can generate more contiguous and complete assemblies from the same library type. However, systematic errors produced by the ONT MinION cannot be fully removed by the R2C2 consensus process or medaka polishing. The assembly we generate does therefore have more mismatches and indel errors than its Illumina counterpart. This ultimately suggests that when limited to a single Tn5 library due to sample constraints, R2C2 can be a valuable addition to an assembly effort but, depending on use case, further polishing with Illumina data might be required to achieve the desired base accuracy.

### Evaluating R2C2 for the sequencing of target-enriched Illumina Tn5 libraries

We tested the ability of R2C2 to evaluate target-enriched Tn5 libraries and benchmark our ability to detect germline variants in the resulting data. To this end, we generated dual-indexed Tn5 libraries from genomic DNA of two cancer cell lines (NCI-H1650 and NCI-H1975) with known mutations in the EGFR gene. We pooled these libraries and enriched the pool for a panel of cancer genes based on the Stanford solid tumor STAMP panel(24) using a Twist Bioscience oligos panel and reagents (Table S2). We performed this enrichment experiment once and without optimization. To compare R2C2 and Illumina MiSeq, we sequenced these enriched Tn5 libraries on 1) a multiplexed Illumina MiSeq 2×300 bp paired end run and 2) on an ONT MinION after R2C2 conversion.

The multiplexed MiSeq run generated 7,430,624 read pairs for the NCI-H1650 library and 1,142,187 read pairs for the NCI-H1975 library. The ONT MinION run generated 3,825,657 R2C2 reads after C3POa processing. Demultiplexing then assigned 2,057,155 (53.7%) R2C2 reads to the NCI-H1650 library and 1,021,758 (26.7%) R2C2 reads to NCI-H1975. Although 537,997 (14.1%) R2C2 reads were not assigned to any sample, only 5.4% of reads were assigned to one of the two combinations of Illumina indexes not included in the pool. The 5.4% single index switching rate implies that, assuming index switches occur independently, only 0.29% (5.4%*5.4%) or 3 out of 1,000 reads would be misassigned to the wrong sample in our dual indexed library.

After demultiplexing we compared the insert length and target enrichment across samples and methods. We did so by merging the Illumina MiSeq read pairs using bbmerge(25). As with the ChIP-seq experiment, R2C2 data showed longer insert lengths than the Illumina MiSeq, with the R2C2 insert length more closely resembling the actual length of the input library (Fig. 5A, D, and S1). We aligned the reads of different samples and methods to the human genome using the short-read preset of minimap2 and determined the percentage of reads overlapped with a target region and the coverage for each region. For NCI-H1650, 15.8% of R2C2 reads and 14.4% of Illumina MiSeq reads overlapped with a target region producing a median coverage of 128 (5th percentile: 28; 95th percentile: 310) for R2C2 and 558 (5th percentile: 134; 95th percentile: 1220) for Illumina MiSeq. For NCI-H1975, 18.5% of R2C2 reads and 16.8% of Illumina MiSeq reads overlapped with a target region with a median coverage of 69 (5th percentile: 13; 95th percentile: 166) for R2C2 and 110 (5th percentile: 23; 95th percentile: 225) for Illumina MiSeq. The per-base coverage of R2C2 and Illumina MiSeq datasets was very well correlated within samples with NCI-H1650 showing a Pearson’s r=0.91 and NCI-H1975 showing a Pearson’s r=0.89 (Fig. 5B and E).

**Fig. 5.**
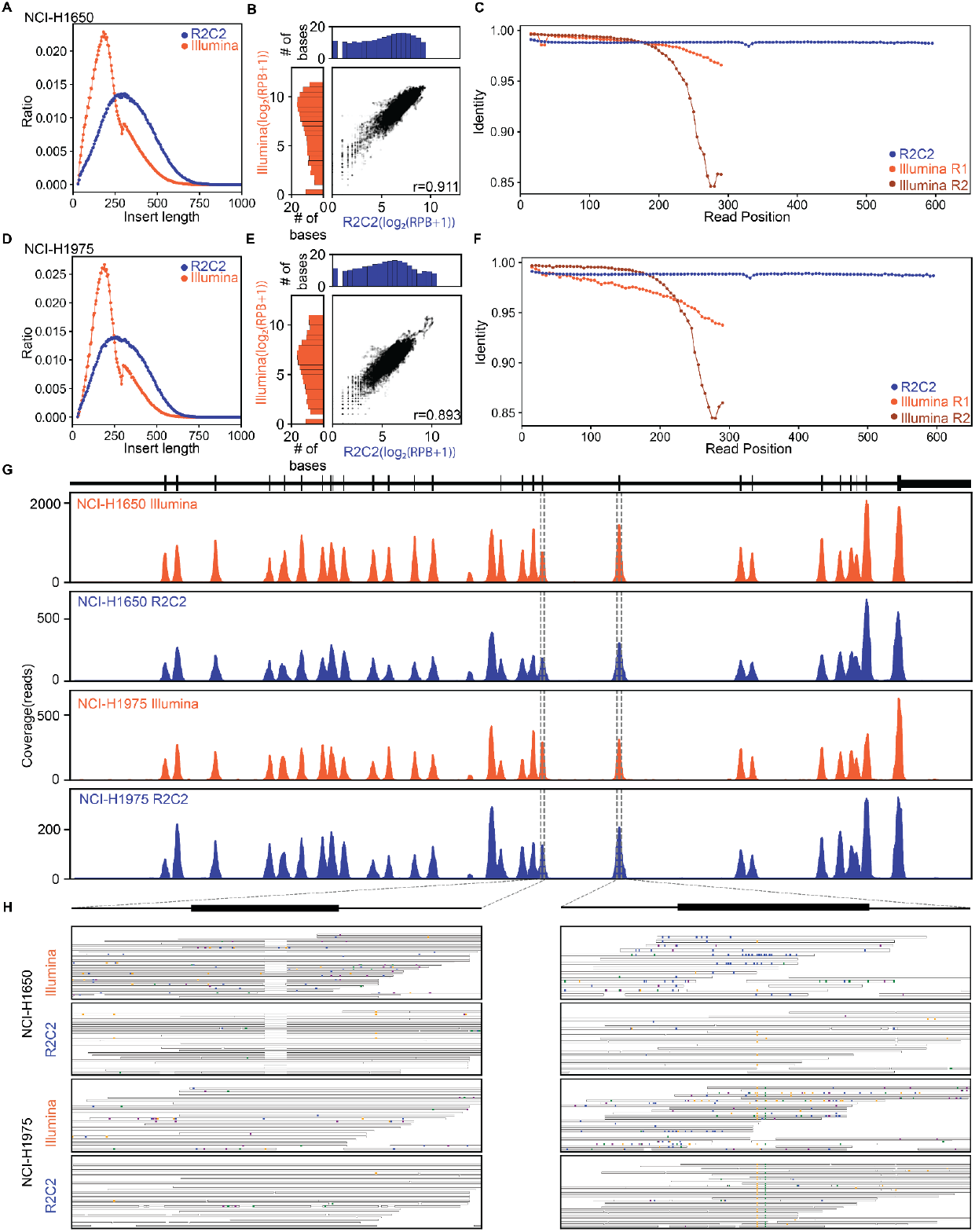
Evaluating target-enriched Tn5 libraries with R2C2. A and D) Inserts length of library molecules sequenced by Illumina or R2C2 approaches. B and E) Comparison of per-base coverage in Illumina and R2C2 datasets. Marginal distributions are log2 normalized. C and F) Alignment based read position dependent accuracy shown for the indicated sequencing reads and methods. Sequencing coverage plot of the target-enriched Tn5 libraries for R2C2 and Illumina results at chromosome 7:55,134,584-55,211,629 which covers a part of the EGFR gene. Top panel shows the annotation of one EGFR isoform. The x axis of the coverage plot is the base pair position and the y axis is the total number of reads at each position. The dotted lines indicate zoomed-in views of exons that contain the 15 bps deletion in NCI-H1650 (left) and the C to T and T to G point mutations in NCI-H1975 (right). Both samples’ Illumina reads and the R2C2 read alignments of the selected regions are shown. The mismatches are colored based on the read base (A - orange; T - green; C - blue; G - purple).

Next, we used the read alignments to determine per-base accuracy levels for all samples and method combinations. The NCI-H1975 sample - which also produced fewer reads than expected on the Illumina MiSeq - produced reads at lower than expected accuracy. Read alignments suggested that the average per-base accuracy for read 1 and read 2 in NCI-1975 were 96.81% and 98.26% compared to 98.37% and 97.88% for NCI-H1650. As expected the per-base accuracy was highly position dependent and declined with increasing sequencing cycle number (Fig. 5C and F). Furthermore, the actual accuracy of the MiSeq reads is likely even lower due to alignments not being extended once the read and genome are too dissimilar. The accuracy of R2C2 reads in both NCI-H1975 and NCI-H1650 were similar and stable throughout the reads at 98.40% and 98.28%, meaning that, in this case, the R2C2 reads had a higher per-base accuracy than the combined MiSeq reads.

Visualizing Illumina MiSeq and the R2C2 read alignments showed that both methods successfully enriched for (Fig 5G) and detected the 15 base pair heterozygous deletion in the EGFR gene in the NCI-H1650 cell line and the C to T heterozygous variants in the EGFR gene in the NCI-H1975 cell line (Fig. 5H). To systematically evaluate the germline variant detection ability of Illumina MiSeq and R2C2 reads, we used Deepvariant(26) for calling germline variants based on the Illumina MiSeq data and Pepper-DeepVariant(27), a variant caller designed for nanopore datasets, for calling germline variants in the R2C2 sequencing results. Because of the poor sequencing performance of the Illumina MiSeq for the NCI-H1975 library, we only performed this analysis on NCI-H1650. For NCI-H1650, Illumina/Deepvariant detected 119 variants in the enriched genomic regions when using a QUAL cut-off of >=33.3. R2C2/Pepper-Deepvariant detected 122 variants in the enriched genomic regions when using a QUAL score >= 3.8 including 117 of the 119 Illumina/Deepvariant calls. When we used Illumina/Deepvariant variants as ground truth, the R2C2/Pepper-Deepvariant method achieved 98.3% recall and 95.9% precision.

Overall, this showed that R2C2 can accurately quantify what percentage of molecules in an enriched Tn5 Illumina library overlap with a target region. Despite showing longer insert lengths than the Illumina MiSeq dataset, the R2C2 dataset showed per-base coverage that was highly correlated with the Illumina MiSeq data. Interestingly in this experiment, R2C2 actually showed a higher per-base accuracy than the Illumina MiSeq. However, due to the remaining error in the R2C2 data likely not being random, variants called based on Illumina MiSeq and R2C2 data are very similar yet not identical, with R2C2 data showing some likely false positive calls. This highlights the persisting limitation of even error-corrected ONT data where increasing per-base accuracy is of limited utility if the remaining errors are systematic in nature.

### Real-Time Analysis of Illumina library metrics using PLNK

To enable the rapid evaluation of Illumina sequencing library metrics by R2C2, we created the computational pipeline PLNK (<u>P</u>rocessing <u>L</u>ive <u>N</u>anopore Experiments). PLNK controls real-time basecalling, raw read processing into R2C2 consensus reads, demultiplexing of R2C2 reads, and the alignment of demultiplexed R2C2 reads to a genome. Based on the resulting alignments and the user defined regions of interest, PLNK then determines the on-target percentage and resulting target coverage for each demultiplexed sample. PLNK runs alongside a MinION sequencing run, tracking the creation of new fast5 files and processing them individually in the order they are generated. To do this, PLNK controls several external tools: guppy5 for basecalling, C3POa for R2C2 consensus generation, a separate python script for demultiplexing (based on splint sequences and Illumina indexes), and mappy (minimap2 python library) for aligning reads to a provided genome (Fig. 6A).

**Fig. 6:**
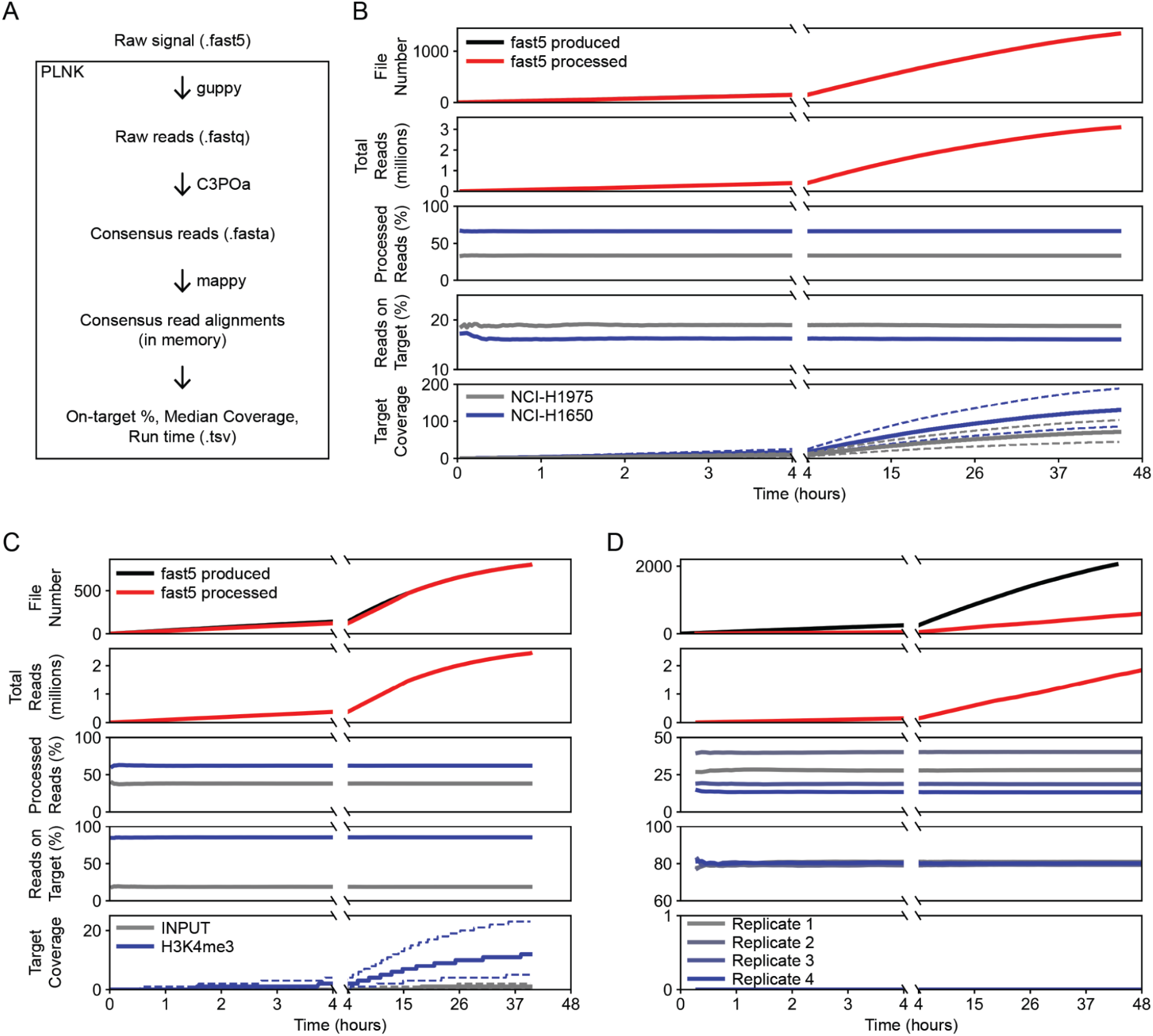
Real-time characterization of Illumina sequencing libraries. A) Diagram of PLNK functionality, fast5 files processed in the order they are produced. PLNK controls guppy5 for basecalling, C3POa for consensus calling, mappy for alignment, and calculates metrics based on those alignments. B-D) Simulation of real-time analysis for enriched Tn5 (B), ChIP-seq (C), and RNA-seq (D) libraries. For each timepoint, panels from top to bottom show 1) The number of fast5 files are produced and processed. 2) The number of demultiplexed reads produced by guppy5/C3POa/demultiplexing. 3) The percentage of reads associated with each library in the sequenced pool. 4) The percent of reads overlapping with target regions 5) The median read coverage of bases in the target regions.

To test whether our pipeline could keep up with ONT MinION data generation and provide real-time analysis, we simulated ONT MinION runs using fast5 files from previously completed sequencing experiments, our Tn5, ChIP-seq and RNA-seq data. We used the fast5 files’ metadata to determine the time intervals at which files were generated by the MinKnow software and copied the fast5 files to a new output directory at those intervals. We then started PLNK to monitor the generation and control the processing of fast5 files in this new output directory. First, we simulated the real-time analysis of the target-enriched Tn5 data. Using a desktop computer and limiting PLNK to the use of eight CPU threads and two Nvidia RTX2070 GPUs, the pipeline processed sequencing data at the same rate a single MinION produced fast5 files. Importantly, both the library composition (percentage of demultiplexed reads assigned to either sample (NCI-H1650 and NCI-1975)) as well as the percentage of reads on-target stabilized after less than an hour and agreed very well with the numbers generated from the whole dataset (Fig. 6B).

When we simulated the analysis of ChIP-seq and RNA-seq experiments, PLNK kept up with ChIP-seq but not with the RNA-seq experiment. Since the RNA-seq experiment produced the largest amount of data in the study, this was not unexpected. In both cases, however, library composition and on-target percent both stabilized within the first hour of sequencing and reflected the number derived from the complete dataset. This means that the library composition and quality of target-enriched Tn5 libraries (as measured by reads overlapping target areas), ChIP-seq libraries (as measured by reads overlapping with peak areas, promoters, or gene bodies - depending on targeted histone mark) and RNA-seq libraries (as measured by reads overlapping with exons) can be determined with minimal sequencing time after which a ONT MinION flow cell can be flushed, stored, and reused.

Overall this suggests that PLNK can be used to evaluate Illumina library metrics in real-time. The bottleneck for analysis in our desktop computer setup seemed to be the guppy5-based basecalling using the slower yet most accurate “sup” basecalling setting. While we could use a faster, less accurate setting to keep up with even the fastest data producing experiments, using the most accurate model means the data can be used for in-depth analysis once the run has completed and PLNK has processed all the files, without the need to re-basecall the raw data.

## Discussion

The capabilities of the dominant Illumina sequencing technology - producing massive numbers of very short reads at moderate accuracy - have shaped the development of sequencing based assays more than any other single factor. While long-read sequencers by Pacific Biosciences (PacBio) and ONT have now clearly superseded Illumina instruments as the gold standard technology for genome assembly, there are still hundreds of assays adapted for very short Illumina reads. These assays are highly diverse and require different levels of read numbers and accuracy and many, like standard RNA-seq, ChIP-seq or targeted sequencing of PCR amplified genomic DNA are unlikely to ever take advantage of the raw read length ONT and PacBio sequencers provide. However, there have been several studies to take advantage of long-read sequencing instruments in sequencing shorter molecules. Some assays [OCEAN, MAS-Iso-Seq] work by either concatenating (8, 28) or otherwise preparing(29) short molecules for sequencing on the PacBio or ONT instrument. While these assays can generate more short reads, they either have to contend with the high cost of the PacBio Sequel IIe sequencer, or the low per-base accuracy of raw ONT reads which even with the latest guppy5 algorithm is only 96% in our hands.

Taking inspiration from the highly accurate but throughput-limited PacBio IsoSeq and HiFi workflows, circularizing-based [R2C2(12), INC-seq(30), HiFRe(31)] methods have been developed to trade throughput for accuracy on ONT MinION and PromethION sequencers. Using a modified R2C2 method we present here, we show that we can convert any Illumina sequencing library with double-stranded adapters - PCR-free “crocodile adapter”-style libraries will not work - into an R2C2 library that takes full advantage of the ONT MinION’s throughput. Overall, these R2C2 libraries produced not only more accurate reads but also a higher number of total reads than regular ONT 1D libraries of the same short-insert Illumina libraries. In fact, throughput and accuracy of R2C2 was comparable to an Illumina MiSeq 2×300 bp run. By generating up to 8.99 million reads (8.1 million demultiplexed) with a per-base accuracy of 98.68% (Illumina MiSeq read 1: 99.47%; read 2: 98.57%) from a single ONT MinION flow cell, this approach can compete with the Illumina MiSeq on accuracy and cost - even without taking instrument cost into account.

We have shown the capabilities and limitations of this approach here by evaluating the conversion of RNA-seq, ChIP-seq, genomic Tn5, and target-enriched genomic Tn5 libraries. The R2C2 data was more than accurate enough to demultiplex Illumina libraries based on their i5 and i7 indexes. Furthermore, RNA-seq data produced with R2C2 were almost entirely interchangeable with data produced by the Illumina MiSeq. Library metrics derived from R2C2 data generated from ChIP-seq and target-enriched Tn5 libraries showed library metrics very similar to those determined from data generated by Illumina sequencers. The notable exceptions to this were insert length distributions of Illumina libraries where R2C2 produced longer insert distributions than Illumina sequencers which are known to prefer shorter molecules enough to affect analysis outcomes(32). The only true limitation of the R2C2 method is rooted in the systematic errors of the ONT sequencing platform that even the R2C2 consensus read approach cannot correct. As a result, even machine learning based polishing (Medaka) or variant calling (Pepper-Deepvariant) tools used in this study couldn’t quite achieve Illumina-style performance for assembly consensus accuracy or germline variant calling, respectively. In either case, it certainly didn’t help that both Medaka and Pepper-Deepvariant had been trained on raw ONT data, not R2C2 consensus reads derived from raw ONT data. Improved consensus tools(33) and consistently improving ONT sequencing accuracy will no doubt also improve R2C2-based performance for these types of analysis in the future.

Beyond simply competing with the Illumina MiSeq, R2C2 could be used for a new group of assays around “medium-length” 600-2000nt reads. Libraries with insert lengths of this size can be size-selected from standard Illumina library preparations. Among many potential applications, this could be applied to standard fragmentation-based RNA-seq libraries to provide more contiguous splicing information for very long transcripts (>15kb) where full-length cDNA based approaches fail.

One of the unique strengths of ONT-based sequencing methods is that, beyond the standard approach of analyzing sequencing runs once they are completed, many library metrics can be derived in real-time. This is starting to get exploited in clinical and metagenomics assays with tools like SURPIrt(34) or with more powerful tools like MinoTour(35). The PLNK tool we developed here controls basecalling by the guppy5 basecaller, C3POa processing, mappy-based alignment and on-target estimation for enrichment analysis. Using this script, we showed that key metrics of RNA-seq, ChIP-seq and enriched Tn5 libraries can be evaluated in under 1 hour of sequencing. This can accelerate quality control and enable the reuse of MinION flow cells, thereby reducing sequencing cost.

In summary, we have shown that, using R2C2, the ONT MinION can - with few limitations - be used as an accurate short-read sequencer with several advantages over dedicated short-read sequencers. Because the ONT MinION comes with minimal instrument cost, R2C2 allows standard short-read genomic assays to be performed in any lab immediately after a library is produced. The presence of a capable short read sequencer in most molecular biology or clinical labs could therefore be truly disruptive by eliminating long turnaround times and therefore dramatically accelerating small-scale experiments.

Even if sequencing libraries are ultimately sequenced in a sequencing core facility on an Illumina HiSeq or NovaSeq 6000 to take advantage of the extremely high throughput these instruments provide, R2C2 can be used to rapidly evaluate library pool compositions and metrics before committing to the cost and turnaround time this requires.

## Methods

### Library Preparation

#### RNA-seq

Four RNAseq libraries were prepared with the NEBNext Ultra II Directional RNA Library Prep Kit for Illumina (NEB #E7760) following the manufacturer’s protocol. For each library, 100 ng of polyA selected RNA from the human lung carcinoma cell line A549 (Takara #636141) was used as input. The RNA fragmentation step was performed at 94C for 5 minutes. PCR enrichment of adaptor ligated DNA was performed for 9 cycles using the NEBNext Multiplex Oligos for Illumina (NEB #E7600S) kit to add Illumina dual index sequences. Three libraries were pooled at a 4ng, 2ng, and 1ng before sequencing on an Illumina MiSeq instrument for paired end 2×300 bp sequencing. The same three RNAseq libraries were pooled again at the same ratio for further R2C2 library preparation. For the 1D and R2C2 runs, the fourth RNA-seq library was prepared and added right before ONT library preparation.

#### ChIP-seq

Chromatin immunoprecipitation (ChIP) was performed following the detailed protocol of Ricci et al. with minor modification(36). In brief, approximately 30 developing seeds at the cotyledon stage were used for chromatin extraction. Immediately after harvesting, the tissue was crosslinked as described in the referenced protocol and immediately flash-frozen in liquid nitrogen. To make antibody-coated beads, 25μl Dynabeads Protein A (Thermo Fisher Scientific, 10002D) were washed with ChIP dilution buffer and then incubated with 2μg antibodies (anti-H3K4me3, Millipore-Sigma, 07–473) for at least 3 hours at 4 °C. After the nuclei extraction, the lysed nuclei suspension was sonicated to 200-500 bp on a Diagenode Bioruptor on the high setting for 30 min. Tubes were centrifuged at 12,000g for 5 min. at 4 °C and the supernatant was transferred to new tubes. At this point, 10 μl of ChIP input aliquots were collected. Sonicated chromatin was diluted tenfold in the ChIP dilution buffer to bring the SDS buffer concentration down to 0.1%. The diluted chromatin was incubated with antibody-coated beads at 4 °C overnight, then washed and reverse-crosslinked. The library was prepared in accordance with the referenced protocol.

#### Tn5

Genomic DNA from a *Wolbachia*-containing *Drosophila Melanogaster* cell line was extracted using a lysis-buffer plus SPRI-bead purification. The Tn5 reaction was then performed using 1ul (22ng) of this genomic DNA, 1ul of the loaded Tn5-AR, 1ul of the loaded Tn5-BR, 13 ul of H2O and 4 ul of 5× TAPS-PEG buffer and incubated at 55°C for 8 minutes (Table S1). The Tn5 reaction was inactivated by cooling down to 4°C and the addition of 5 μl of 0.2% sodium dodecyl sulfate then incubated for 10 minutes. 5 ul of the resulting product was nick-translated at 72°C for 5 minutes and further amplified using KAPA Hifi Polymerase (KAPA) using Nextera Index primers with an incubation of 98°C for 30 s, followed by 16 cycles of (98°C for 20 s, 65°C for 15 s, 72°C for 30s) with a final extension at 72°C for 5 min. Before R2C2 conversion, the resulting Tn5 library was size-selected for molecules between 800-1200bp on a 1% low-melt agarose gel.

#### Target-enriched Tn5

The Tn5 library was prepared using genomic DNA from cell lines NCI-H1650 (ATCC CRL-5883D) and NCI-H1975 (ATCC CRL-5908DQ). A total of 100ng genomic DNA of each sample was treated with Tn5 enzyme loaded with Tn5ME-A/R and Tn5ME-B/R. The Tn5 reaction was performed using 1ul of the gDNA, 1ul of the loaded Tn5-AR, 1ul of the loaded Tn5-BR, 13 ul of H2O and 4 ul of 5× TAPS-PEG buffer and incubated at 55°C for 8 minutes. The Tn5 reaction was inactivated by cooling down to 4°C and the addition of 5 μl of 0.2% sodium dodecyl sulfate then incubated for 10 minutes. 5 ul of the resulting product was nick-translated at 72°C for 5 minutes and further amplified using KAPA Hifi Polymerase (KAPA) using Nextera_Primer_B_Universal and Nextera_Primer_A_Universal (Smart-seq2) with an incubation of 98°C for 30 s, followed by 16 cycles of (98°C for 20 s, 65°C for 15 s, 72°C for 30s) with a final extension at 72°C for 5 min.

The resulting Tn5 library was then enriched with Twist fast hybridization reagents and customized oligo panels that were designed based on the Stanford STAMP panel. The hybridization reaction of the panel and the Tn5 libraries was performed using 294ng of NCI-H1975 Tn5 library, 360ng of NCI-H1650 Tn5 library, 8ul of blocking oligo pool [100uM], 8ul of universal blockers, 5ul of blocker solution and 4ul of the custom panel. The mix was dehydrated using SpeedVac and was resuspended in 20ul Fast Hybridization mix at 65C. After the addition of 30 ul of Hybridization Enhancer, the mixture was incubated at 95C for 5 minutes and 60C for 4 hours. After hybridization, the reaction mix was incubated with pre-washed Streptavidin binding beads and washed using the Fast Wash buffer one and Fast Wash buffer two for six times. The Streptavidin beads and the DNA mixture was used directly for reamplification with Universal primers and Equinox Library Amp Mix. The mixture was incubated at 98°C for 45 s, followed by 16 cycles of (98°C for 15 s, 65°C for 30 s, 72°C for 30s) with a final extension at 72°C for 1 min. The final enriched Tn5 library DNA product was cleaned up using SPRI beads at 1.8:1 (Beads:Sample) ratio.

#### R2C2 Conversion

Pooled Illumina libraries were first circularized by Gibson assembly with a DNA splint containing end sequences complementary to ends of Illumina libraries (Table S1). Illumina libraries and DNA splint were mixed at a 1:1 ng ratio using NEBuilder HiFi DNA assembly Master mix (NEB #E2621). Any non-circularized DNA was digested overnight using ExoI, ExoIII, and Lambda exonuclease (all NEB). The reaction was then cleaned up using SPRI beads at a 0.85:1 (Bead:Sample) ratio. The circularized library was then used for an overnight RCA reaction using Phi29 (NEB) with random hexamer primers. The RCA product was debranched with T7 endonuclease (NEB) for 2 hours at 37C then cleaned using a Zymo DNA Clean & Concentrator column-5 (Zymo #D4013). The cleaned RCA product was digested using NEBNext dsDNA Fragmentase (NEB #M0348) following the manufacturer protocol with a 10 minute incubation. For the regular Tn5 library digested RCA product was cleaned using SPRI beads. For all other libraries, the digested RCA product was size selected using a 1% low melt agarose gel: DNA between 2-10 kb was excised from the gel which was then digested using NEB Beta-Agarase. DNA was then cleaned using SPRI beads.

#### ONT sequencing

ONT libraries were prepared from R2C2 DNA or directly from Illumina libraries using the ONT ligation sequencing kit (ONT #SQK-LSK110) following the manufacturer’s protocol then sequenced on an ONT MinION flow cell (R9.4.1). When preparing ONT libraries from Illumina libraries, SPRI bead purifications throughout the protocol were adjusted to accommodate for their short length. Additional library was loaded on the same flow cell after nuclease flush.

#### Illumina sequencing

Library pools were sequenced either on the Illumina MiSeq using 2×300 (RNA-seq and target enriched Tn5 libraries), the Illumina NextSeq500 2×150 (Tn5 library) or the Illumina HiSeq (ChIP-seq)

### Analysis

#### R2C2 and 1D

Raw nanopore sequencing data in the fast5 file format was basecalled using the “sup” setting of guppy5 to generate fastq files. R2C2 raw reads in fastq format were then processed by C3POa (v.2.2.3 - https://github.com/rvolden/C3POa) to generate accurate consensus reads. R2C2 consensus reads and ONT 1D reads were further processed with C3POa (C3POa_postprocessing.py), using the --trim setting and the following p5/p7 adapter sequences:

~~~
>3Prime_adapter
CAAGCAGAAGACGGCATACG
>5Prime_adapter
AATGATACGGCGACCACCGAGATCT
~~~

Custom scripts (available at https://github.com/kschimke/PLNK) were used to demultiplex reads based on the sequences of their DNA splints and Illumina indexes and to trim the rest of the Illumina sequencing adapters.

#### RNA-seq

To determine accuracy levels R2C2, 1D, Illumina MiSeq reads were aligned to the human genome reference (hg38) using minimap2 (v2.18-r1015)(18).

~~~
*minimap2 -ax splice --cs=long --MD –secondary=no*
~~~

Position dependent accuracy was determined after converting sam files with the sam2pairwise tool(37).

Illumina reads were adapter trimmed using cutadapt (v3.2)(38)

~~~
*cutadapt -m 30 -j 50 -a AGATCGGAAGAGC -A AGATCGGAAGAGC*
~~~

Illumina and R2C2 reads were aligned to the human genome (hg38) using STAR and STARlong (v2.7.3a)(16)

~~~
*STAR --quantMode GeneCounts --outSAMattributes NH HI NM MD AS nM jM jI XS*
~~~

To determine insert length, Illumina read pairs were merged using bbmerge (v38.92) with default settings.

#### ChIP-seq

Illumina reads were sub-sampled using a custom script (https://github.com/alexanderkzee/BWN) to match the total reads from the corresponding R2C2 library.

Illumina and R2C2 reads were aligned to the *Glycine Max* genome (Gmax_508_v4.0) using minimap2 (v2.18-r1015)(18).

~~~
*minimap2 -ax sr --cs=long --MD --secondary=no*
~~~

Peaks in H3K4me3 Illumina data were called using MACS2(39)

~~~
*macs2 callpeak -t K4.bam -c INPUT.bam -f BAM -n K4_Illumina --nomodel --extsize 200*
~~~

#### Tn5

R2C2 reads were aligned to the *Drosophila melanogaster* genome (dm6) using minimap2 ((v2.18-r1015)

~~~
*minimap2 -ax sr --cs=long --MD --secondary=no*
~~~

R2C2 reads that didn’t align to the Drosophila genome were then assembled using miniasm

~~~
*minimap2 -x ava-ont [dehosted r2c2 file] [dehosted r2c2 file] > [ava paf file]
miniasm -f [dehosted r2c2 file] [ava paf file] -m 450 -s 250 > [gfa raw assembly]*
~~~

We aligned Illumina reads to the Drosophila melanogaster genome (dm6) using bwa mem(40) under default parameters. We then extracted the sample IDs for reads that did not map to the host genome and extract that set from the raw fastq files.

Illumina reads that didn’t align to the Drosophila genome were then assembled using meraculous, setting the minimum contig depth to 10, expected genome size to 0.013, and using a k-mer of 51 and otherwise default parameters.

#### Target-enriched Tn5

Illumina reads were adapter trimmed using cutadapt (v3.2)

~~~
*cutadapt -m 30 -j 50 -a AGATCGGAAGAGC -A AGATCGGAAGAGC*
~~~

Trimmed Illumina and R2C2 reads were aligned to the human genome (hg38) using minimap2 (v2.18-r1015).

~~~
*minimap2 -ax sr --cs=long --MD –secondary=no*
~~~

Germline variants in Illumina data of NCI-H1650 were called using Deepvariant(26). Germline variant in R2C2 data of NCI-H1650 were called using Pepper-Deepvariant(27)

#### Real-time Analysis with PLNK

RNA-seq, ChIP-seq and Enriched Tn5 MinION runs were simulated by reading the *mtime* metadata entry of fast5 files in the output folder of the completed runs and then calculating the time intervals at which files were created by the MinKNOW software. Files created during the first 48 hours or until the first library reload were then copied into a new folder at those intervals. PLNK (https://github.com/kschimke/PLNK) was started after the simulation and was given key information about the run (splint and Illumina indexes in the format of a sample sheet, target regions in bed format, genome sequence in fasta format) and a config file containing paths to tools used by PLNK.

samtools(41) (v1.11-18-gc17e914) was used extensively during analysis for sam file processing. Python(42), matplotlib(43), numpy(44), and scipy(45) were all used to analyze and visualize the data

## Data availability

All data generated and analyzed for this paper are available as Bioproject PRJNA775962 at the SRA.

## Code availability

All code used for analysis is available at the following github repositories

https://github.com/kschimke/PLNK

https://github.com/alexanderkzee/BWN

https://github.com/rvolden/C3POa as indicated throughout the method section.

## Acknowledgements

We thank the UCSC Paleogenomics Lab sequencing facility for sequencing RNA-seq, Tn5 and Enriched Tn5 libraries. We also want to thank Kishwar Shafin for his support with running Pepper-Deepvariant. We acknowledge funding by the National Institute of General Medical Sciences / National Institutes of Health Grant R35GM133569 (to C. V.) and R35GM128932 (to R.C.-D.). This study was funded with support from the NSF (IOS-1856627) and the United Soybean Board to R.J.S..

## Author contributions

A.Z. led the R2C2 conversion of ChIP-seq libraries and Tn5 libraries, analyzed ChIP-seq data, wrote and edited the manuscript.

D.D. led the generation and R2C2 conversion of target-enriched Tn5 libraries, analyzed the target-enriched Tn5 data, wrote and edited the manuscript.

M.A. optimized the R2C2 method for the conversion of Illumina libraries, led the generation and R2C2 conversion of RNA-seq libraries, analyzed the RNA-seq data, wrote and edited the manuscript.

K.S. led the development of the PLNK real-time analysis tool, performed real-time analysis simulations, wrote and edited the manuscript.

R.C.D. conceptualized and supervised the generation of Tn5 libraries, analyzed the Tn5 library based genome assemblies, and edited the manuscript.

S.R. conceptualized the generation of Tn5 libraries, generated *Wolbachia*-containing *Drosophila melanogaster* cell lines, and edited the manuscript.

X.Z.generated and supervised the Illumina sequencing of ChIP-seq libraries, and edited the manuscript.

R.J.S.designed and supervised the generation of ChIP-seq libraries and data, and edited the manuscript.

C.V.conceptualized and designed the project, supervised the generation of data, supervised Illumina and R2C2 data analysis, performed germline variant calling on target-enriched Tn5 Illumina and R2C2 data, wrote and edited the manuscript

## Supplementary material

**Fig S1.**
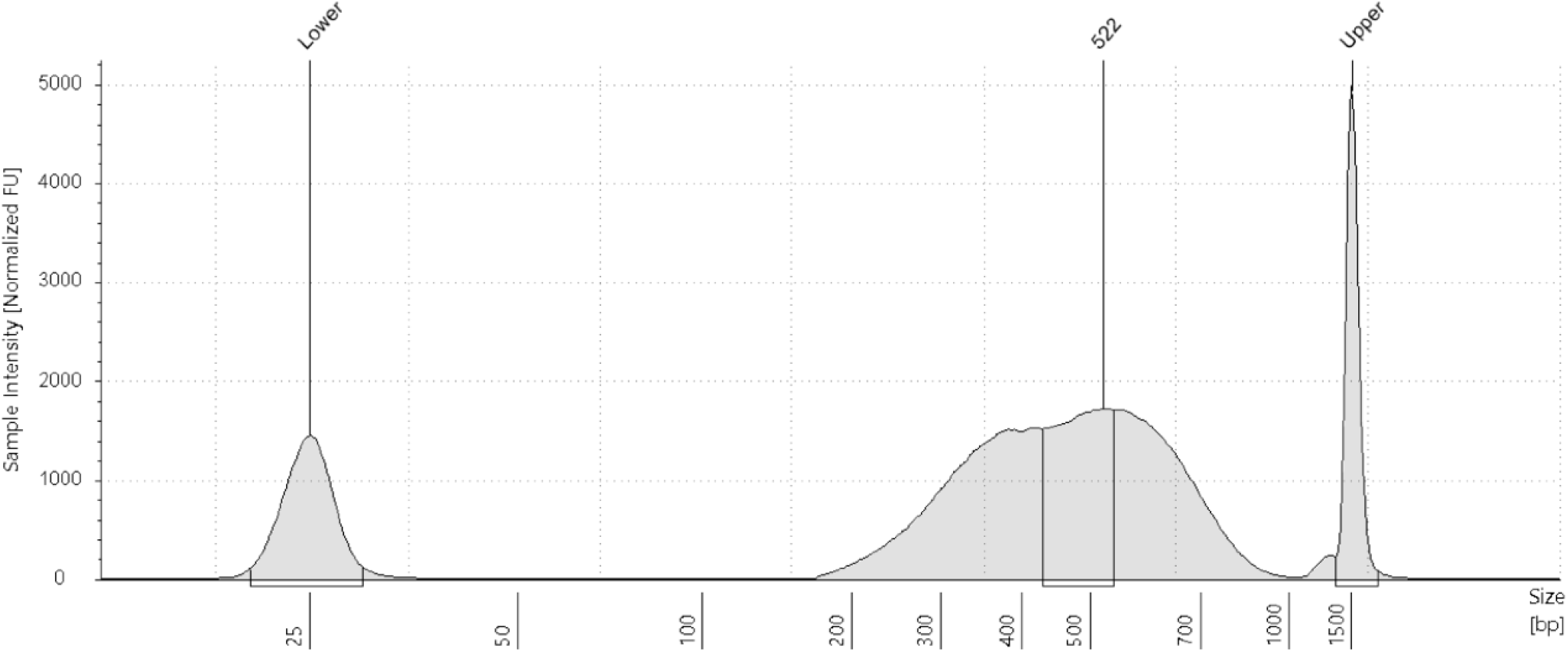
Target-Enriched Tn5 library size. The size of the target-enriched Tn5 library pool as determined by Agilent Tapestation run.

**Table S1:**
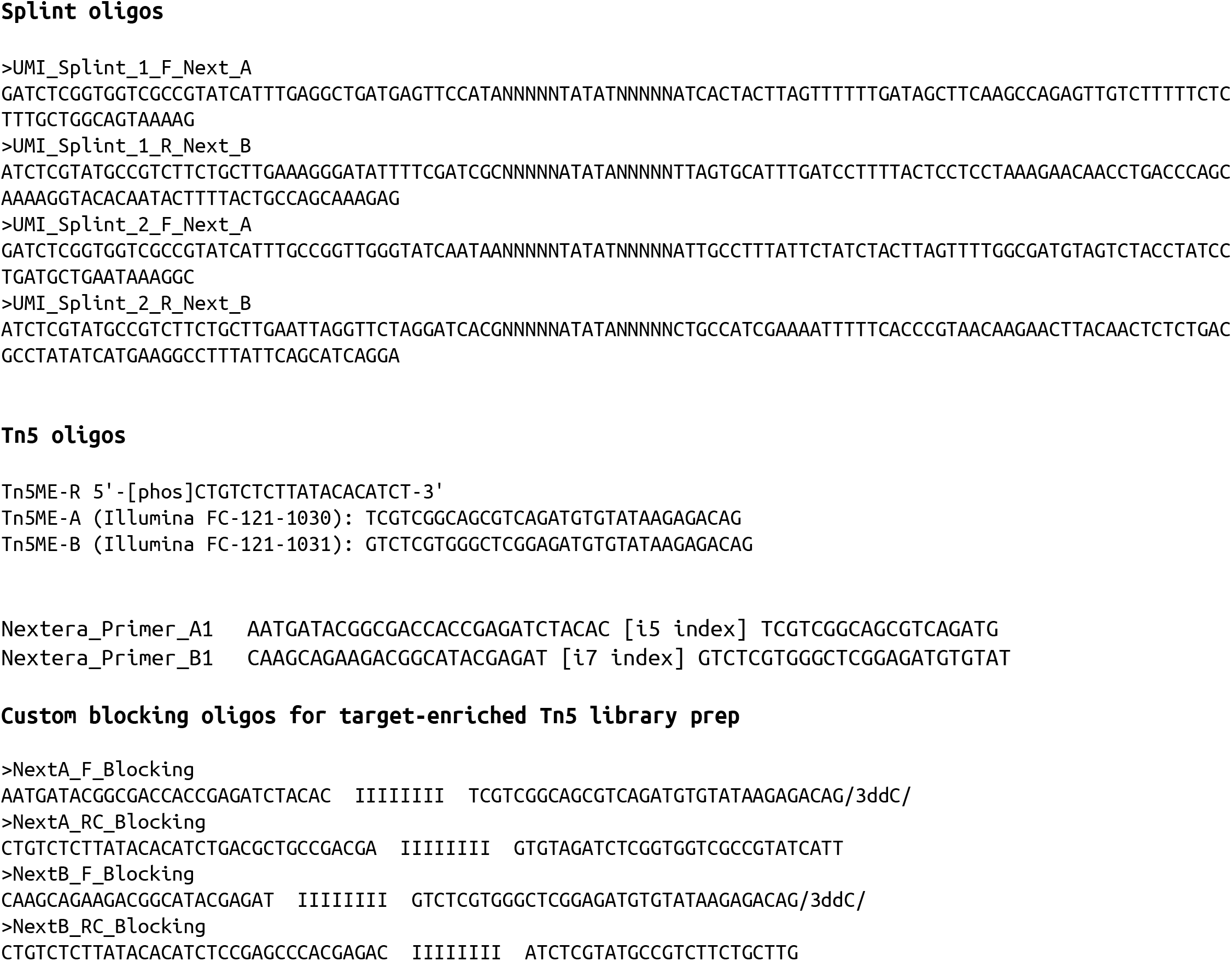
Custom oligos used in the IBWN study.

## Notes

### Competing Interest Statement

Oxford Nanopore Technologies (ONT) has paid for travel and accommodations for C.V. to their London Calling conference twice between 2016 and 2018. In 2020, ONT supplied sequencing reagents to the lab of C.V. for work on the LRGASP consortium - an unrelated project.

### Summary of Updates

In addition to several updates to text, tables, and figures, we added regular ONT ligation prep sequencing results for Illumina RNA-seq libraries for comparison.

